# Delayed accumulation of inhibitory input explains gamma frequency variation with changing contrast in an Inhibition Stabilized Network

**DOI:** 10.1101/2024.07.04.602155

**Authors:** R Krishnakumaran, Abhimanyu Pavuluri, Supratim Ray

## Abstract

Gamma rhythm (30-70 Hz), thought to represent the push-pull activity of excitatory and inhibitory population, can be induced by presenting achromatic gratings in the primary visual cortex (V1) and is sensitive to stimulus properties such as size and contrast. In addition, gamma occurs in short bursts, and shows a “frequency-falloff” effect where its peak frequency is high after stimulus onset and slowly decreases to a steady state. Recently, these size-contrast properties and temporal characteristics were replicated in a self-oscillating Wilson-Cowan (WC) model operating as an Inhibition stabilized network (ISN), stimulated by Ornstein-Uhlenbeck (OU)-type inputs. In particular, frequency-falloff was explained by delayed and slowly accumulated inputs arriving at local inhibitory populations. We hypothesized that if the stimulus is preceded by another higher contrast stimulus, frequency-falloff could be abolished or reversed, since the excessive inhibition will now take more time to dissipate. We presented gratings at different contrasts consecutively to two female monkeys while recording gamma using microelectrode arrays in V1 and confirmed this prediction. Further, this model also replicated a characteristic pattern of gamma frequency modulation to counter-phasing stimuli as reported previously. Thus, the ISN model with delayed surround input replicates gamma frequency responses to time-varying contrasts.

**Significance statement:** Gamma rhythms represent sustained push-pull dynamics between excitatory and inhibitory populations during visual stimulation. Gamma power and centre frequency varies depending on stimulus features, and onset of stimulus produces a “frequency-fall” trend where onset frequency is higher and subsequently plateaus to a lower value. In an earlier work, we argued, using a noisy rate-model of V1, that a delayed onset of inhibition-drive from the surround populations produced the gamma ‘frequency-falloff’. We tested a key prediction of this hypothesis that the frequency-falloff can be abolished or reversed if the stimulus is preceded by a higher contrast stimulus, and confirmed the same by recording from primate primary visual cortex while presenting multiple stimuli consecutively at varying contrasts.

## Introduction

Gamma rhythm (30-70 Hz) can be induced in the primary visual cortex (V1) by presentation of suitable achromatic gratings or plain hues, and represents the intrinsic push-pull activity of excitatory and recurrently connected inhibitory populations. Such gamma generation has been demonstrated in network models of V1 (Buzsáki and Wang, 2012; Chariker et al., 2018; Zachariou et al., 2021), suggesting gamma rhythm characteristics are tightly linked to the recurrent circuit within V1. Gamma properties tend to reflect possible changes in cortical circuitry due to ageing (Murty et al., 2020) and mild cognitive impairment (MCI; Murty et al. 2021).

Gamma rhythm exhibits differences in power and peak frequency depending on the properties of stimulus presented. For instance, gamma frequency is higher during presentation of higher contrast stimuli (Ray and Maunsell, 2010), while gamma power increases and frequency decreases when larger size stimuli are presented (Gieselmann and Thiele, 2008; Ray and Maunsell, 2011; Peter et al., 2019). These properties of V1 gamma had been replicated in a Wilson-Cowan (WC)-type model, which we refer to as the JS model, that generate gamma as limit cycles (Jadi and Sejnowski, 2014; Shirhatti et al., 2022). Shirhatti and colleagues have previously shown that the existence of bifurcation in the JS model emulates the attenuation of gamma rhythm induced by a fullscreen achromatic grating when a small discontinuity is introduced near the receptive field (RF) centre (Shirhatti et al., 2022). Furthermore, gamma rhythms exhibit specific temporal characteristics such as non-sinusoidal waveform shape (Krishnakumaran et al., 2022), and bursty nature, which have also been replicated in the JS model (Krishnakumaran and Ray, 2023). Gamma bursts with realistic burst length distributions could be replicated in the JS model by supplying noisy inputs of Ornstein-Uhlenbeck (OU) type, obtained by low-pass filtering of Poisson distributed inputs. Furthermore, gamma rhythms exhibit a “frequency-falloff”, where the gamma frequency is high near stimulus onset, and slowly reduces to a lower steady-state value, which is visible over the initial 300-400 ms in time-frequency (TF) spectra (For examples, see TF spectra in Fig. 1 of (Xing et al., 2012), Fig. 5 of (Peter et al., 2019) for LFP in macaques, Fig. 4a in (Murty et al., 2020) for EEG in humans and Fig. 3 of (Perry et al., 2020) for MEG in humans; The falloff is more visible in macaque LFP owing to better signal-to-noise ratio of LFP). Replicating the frequency-falloff in the JS model required the cut-off of filter be lower for the input drive to the inhibitory population than for the excitatory population (Krishnakumaran and Ray, 2023). This suggested a slower accumulation of inhibition, analogous to the temporal integration of inhibitory drive coming from surround populations over unmyelinated lateral connections arriving at different delays (Angelucci et al., 2017), and the differential synaptic dynamics of excitation and inhibition.

**Figure 1:**
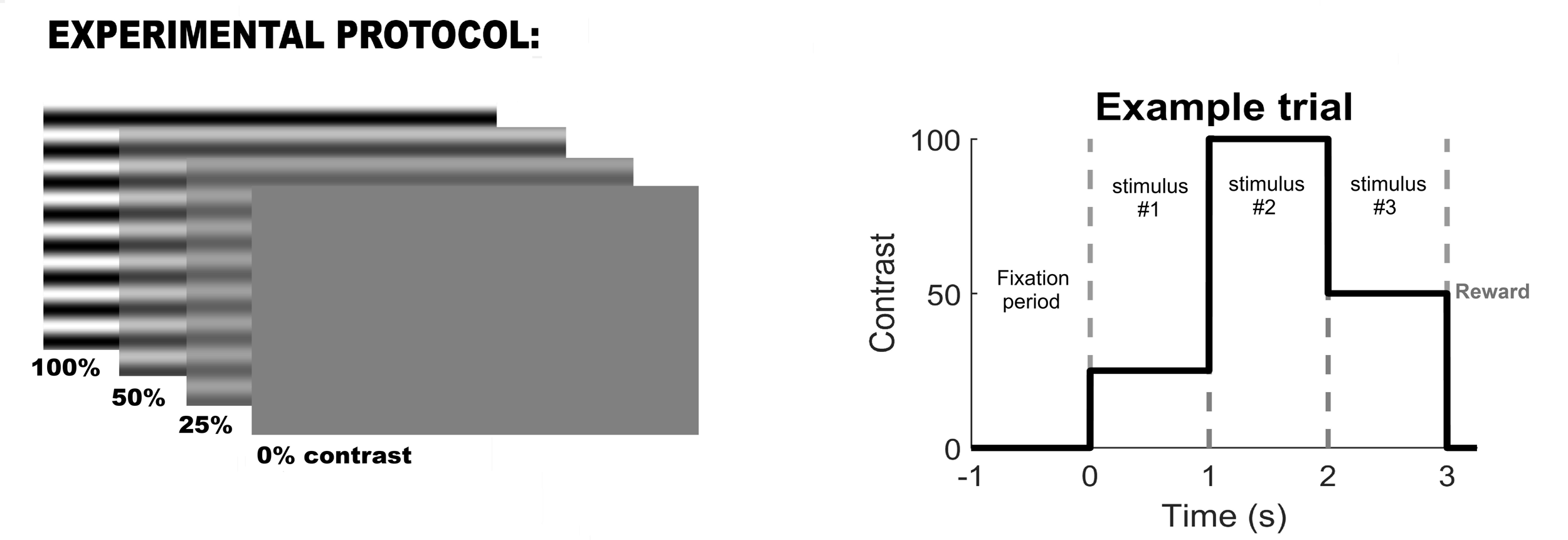
Summary of experimental paradigm and data analysis. Illustration of stimulus presentation during an example trial of passive fixation task. A fullscreen achromatic grating is presented in 3 contrasts and a blank grey screen is presented during the fixation period and as 0% contrast stimulus. A single trial of the passive fixation task consists of a fixation period followed by 3 consecutive contrast presentations without interstimulus intervals. Maintaining fixation at the centre of the screen throughout the fixation period and the stimulus presentations results in a reward, whereas breaking it causes the trial to end without reward.

As inhibition is slower in accumulation and decay in our model, we predicted that a prior build-up of inhibition, caused by presenting a higher contrast before a lower contrast, would either reverse or suppress the trend of the frequency-falloff for the lower contrast stimulus. In the current study, we explore this effect of contrast adaptation on the frequency-falloff of gamma rhythm by presenting fullscreen gratings of different contrasts consecutively with no inter-stimulus interval. Using time-frequency spectral analysis, we traced the frequency of gamma bursts at different times from stimulus onset, and compared the frequency transient for a stimulus, under unadapted and adapted conditions. Further, we tested whether the noisy JS model replicates a characteristic pattern of gamma frequency modulation in response to slow time-varying contrast stimuli as reported previously (Ray and Maunsell, 2010).

## Materials and Methods

### Ethics Statement

All the experiments on monkeys were conducted in adherence to guidelines approved by the Institutional Animal Ethics Committee of the Indian Institute of Science-Bangalore and the Committee for the Purpose of Control and Supervision of Experiments on Animals (CPCSEA).

### Animal Preparation and Training

Two adult female monkeys (Macaca radiata; M1: 13 y, ∼3.3 kg and M2: 15 y, ∼5.6 kg) were used. Each monkey was surgically implanted with a titanium headpost over the anterior/frontal region of the skull under general anesthesia. Once recovered, the monkey was trained on a visual passive fixation task. After the monkey learnt to maintain fixation for an adequate duration, a second surgery was performed under general anesthesia to insert a microelectrode array (Utah array, 81 active platinum microelectrodes in M1 and 48 electrodes in M2, 1 mm long each, 400 µm interelectrode distance; Blackrock Microsystems) in the primary visual cortex area V1 (right hemisphere, centered ∼10–12 mm rostral from the occipital ridge and ∼10–12 mm lateral from the midline, with location varying slightly in the two monkeys). In M2, a second microelectrode array was placed in V4 in the same surgery, whose signals were not utilized in the current study. The RFs of the recorded V1 neurons were located in the lower left quadrant of the visual space with respect to fixation (at an eccentricity of ∼3°–4.5° in M1 and ∼1.5°–3° in M2). Following a period of post-surgery care and monitored recovery of at least 2 weeks, the monkeys performed the experimental task regularly while microelectrode data were recorded.

### Data Acquisition

Raw signals from microelectrodes were recorded using the 128-channel Cerebus neural signal processor (Blackrock Microsystems). The signals were filtered online between 0.3 Hz and 500 Hz (Butterworth filters; first-order analog and fourth-order digital respectively) to get the LFP data, which were then recorded at a sampling rate of 2 kHz and a resolution of 16-bits. This LFP timeseries was used directly in our analyses.

### Experimental setup and behavior

During the experiment, the monkey’s head was held still by the headpost as it sat in a monkey chair and viewed a monitor (BenQ XL2411, LCD, 1,280 × 720 resolution, 100 Hz refresh rate) placed ∼50 cm from its eyes. A Faraday enclosure housed the subject and the display setup with a dedicated ground for isolation from external electrical noise. The monitor was calibrated and gamma-corrected using i1Display Pro (x-rite PANTONE). Mean luminance was set to 60 cd/m2 at the monitor surface and gamma was set to unity for each of the three primaries.

The monkeys were subjected to a passive fixation task, in which they were rewarded with juice for fixating at a small dot of 0.05°–0.10° radius at the center of the screen throughout the trial (4 s duration; fixation spot was displayed throughout). Each trial began with fixation period of 1 s duration, followed by consecutive presentations of an achromatic grating at three (sometimes two for M1) different contrasts (details on the grating in next section - *Stimuli*), 1 s each with an interstimulus interval of 0 s, as depicted in Figure 1. The order of different contrasts in each trial was randomly chosen. Juice was awarded only if fixation was maintained within 2° from the fixation point. Trials in which fixation was broken were not considered in our analyses. Eye position data was recorded using the ETL-200 Primate Eye Tracking System (ISCAN) which gave horizontal and vertical coordinates. During the task, monitoring of eye position data, control of task flow, generation and randomization of stimuli were performed using a custom software on macOS. Although fixation was required to be within 2° from the fixation spot, the monkeys were able to hold their fixation well within these limits during the task with a standard deviation of less than 0.4° across all sessions for both monkeys. These small eye movements are unlikely to affect our results since full-screen stimuli were used.

### Stimuli

The set of stimuli comprised fullscreen static achromatic grating of different contrasts (Figure 1). The achromatic grating was at an orientation of 90° for both monkeys and had a spatial frequency of 4 cpd for M1 and 2 cpd for M2. These grating parameters were chosen to induce strong fast gamma and reduced slow gamma (Murty et al., 2018). These full-screen stimuli subtended a visual angle of ∼56° in the horizontal direction and ∼33° in the vertical direction. The gratings were presented at 3 different contrasts in M1 (25%, 50% and 100%) and 4 different contrasts in M2 (0%, 25%, 50% and 100%; 0% contrast produces the same blank grey screen as during fixation period). An additional 0% contrast was included in M2 to consider unadapted conditions occurring later in a trial whereas in M1, unadapted conditions for a stimulus would refer to trials where a given contrast was the first to be displayed after fixation period. The results did not differ between unadapted conditions positioned at fixation period and later in trial in M2, and so the trials are pooled together in analyses. Adapted condition refers to the presentation of a stimulus, immediately preceded by a stimulus with non-zero contrast.

### Electrode selection

As with our previous reports (Krishnakumaran et al., 2022; Krishnakumaran and Ray, 2023), we considered only those electrodes for analysis that had robust estimates of the RFs which were estimated by presenting small sinusoidal gratings at different eccentricities from the central fixation point. These criteria were determined by a RF mapping protocol that was run across multiple days (Dubey and Ray, 2019; Krishnakumaran and Ray, 2023). Further, high artifact electrodes were identified using the impedances measured on the days of recording and discarded. This procedure yielded 77 and 32 electrodes from M1 and M2 respectively.

### Data Analysis

We discarded stimulus presentations with excessive artifacts for each session (<5.9% of presentations of each stimulus in M1 and M2). The resultant number of useable trials for each contrast pair is given in the supplementary Figure S1 for M1 and Figure S2 for M2. For the low contrast condition (50 %) we present in the *Results* section, the number of useable trials were 93 trials in M1 and 87 trials in M2 for each unadapted presentations of 50 % contrast and 86 trials in M1 and 35 trials in M2 for the 100% contrast-adapted. The trials in the unadapted case in M2 were more in number than the adapted case since this set included all trials where the non-zero contrast stimulus was presented immediately after the fixation period, and other trials where it was presented later in the stimulus sequence but preceded by a 0 % contrast stimulus. Furthermore, the software used to generate our trials as pseudo-randomly generated sequences of contrasts used a blocked design where different contrasts (25%, 50% and 100% for M1 and 0%, 25%, 50% and 100% for M2) were chosen pseudo-randomly presented without replacement and the next block started only once all contrasts were shown. Therefore, two consecutive stimuli with the same contrast could occur only when one block finished with a particular contrast and the next block started in the same trial with the same contrast, which happened rarely. This gave rise to the very low number of trials with the same contrast presented consecutively (or equivalently a single contrast presented for 2 s), as shown in the diagonal-occupying plots of Figures S1 and S2.

### Time-Frequency analysis of LFP and Gamma peak-frequency estimation

Time-frequency (TF) spectra were computed using Multi-tape method (MT) using the Chronux toolbox (Bokil et al., 2010), with moving windows of 0.4 s and 3 tapers. This gave us a frequency resolution of 2.5 Hz, which was needed to trace the gamma peak-frequency falloff. To compute the electrode averaged change in TF spectral power from baseline, for each stimulus, the TF spectrum from each trial corresponding to the presentation of the stimulus from each electrode were averaged. Then the baseline power at each frequency was computed by averaging the power at the respective frequency over the baseline period interval between - 750 ms to 0 ms from stimulus onset. This baseline for each frequency was subtracted (on a log scale) from the averaged TF spectrum at every timepoint at the frequency. The peak-frequency time-series of gamma was computed for each stimulus as the frequency within 20 to 80 Hz range with the maximum power in the electrode averaged change in TF spectrum at each timepoint.

### Experimental Design and Statistical Analysis

To estimate gamma peak-frequency trace at each time-point, the peak-frequencies estimated from all the electrodes at the respective timepoint were pooled and the median of the resulting distribution was computed and plotted in Figure 2. The standard error (SE) was estimated by bootstrapping over N iterations (where N is the number of electrodes). This involved random sampling with replacement of the data N times and estimating their median in each iteration, which resulted in N medians, whose standard deviation was taken as the SE.

**Figure 2:**
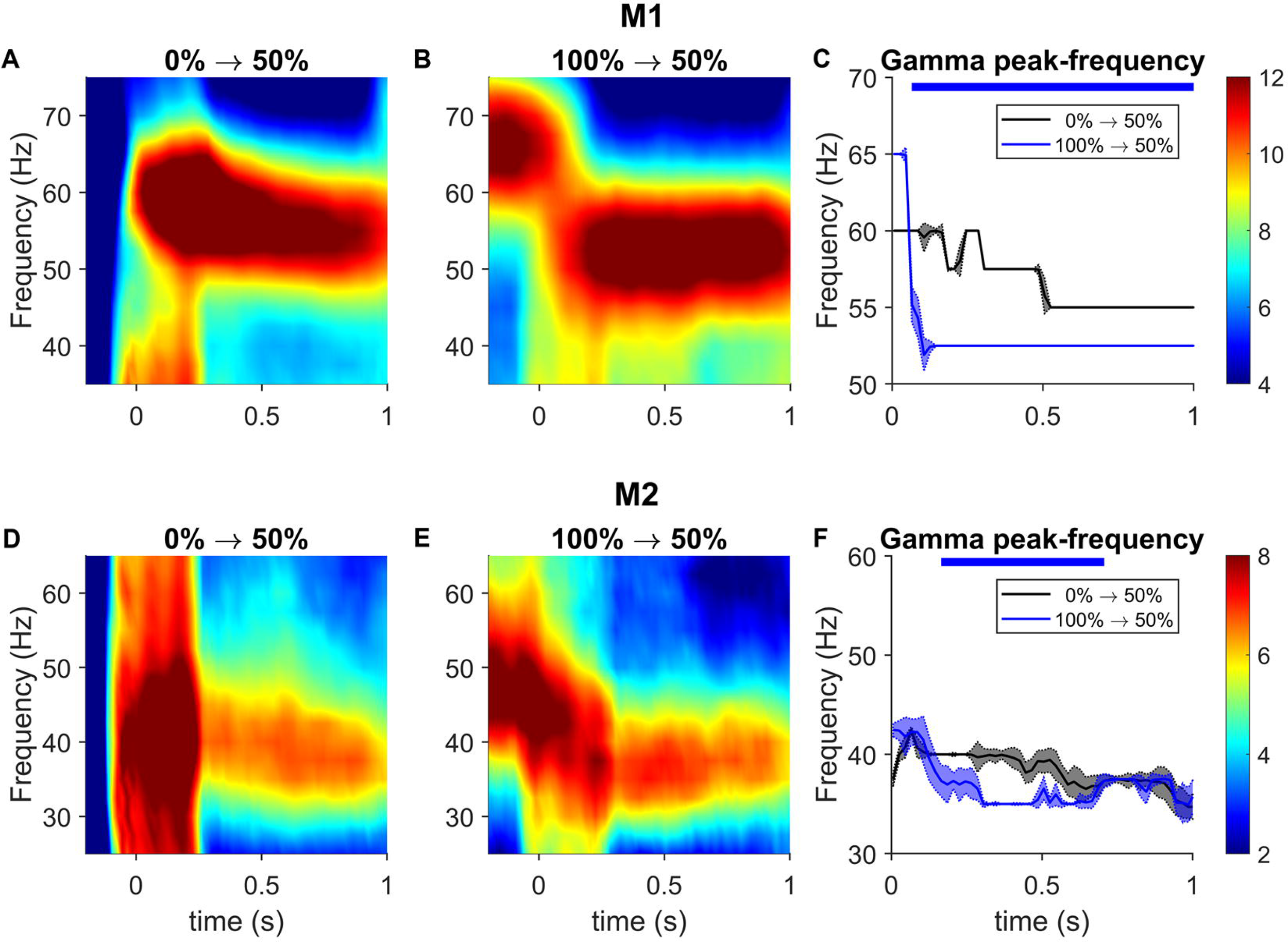
Effect of contrast adaptation on gamma in time-frequency spectrum. (A) Change in TF spectrum of LFP from M1 from unadapted trials where 50% contrast was preceded by blank grey screen. The 0 s mark corresponds to the onset of 50% contrast. (B) Same as A but from trials with 50% contrast following a 100% contrast. (C) Peak frequency of gamma band for each condition. Solid line traces the median peak-frequency of gamma across electrodes. The width of the line at each timepoint represents the standard error (SE), computed using bootstrapping. Black and blue traces indicate gamma-band peak-frequency traces corresponding to change in TF spectra in (A) and (B) respectively. Horizontal blue bars at the top edge mark the timepoints where the difference between adapted and unadapted peak-frequencies is statistically significant (p<0.01). (D-F) LFP spectra and gamma frequency analysis from M2.

To test any difference in temporal trend in gamma frequency between adapted and unadapted conditions, the peak-frequencies were subject to a paired tailed sign-test. First the difference in peak-frequencies recorded in a single electrode between adapted and unadapted conditions was taken at each time-point. This resulted in a difference in frequency time-series for each electrode. This trace was binned into non-overlapping consecutive time windows of 10 ms starting from 0 ms. In each time-window, the difference in frequency values between adapted and unadapted condition across all electrodes were pooled and subject to a right-tailed sign-test. A p-value < 0.01 obtained for a time-window rejects the null hypothesis that the median difference in frequency between adapted and unadapted conditions is 0 within the time-window, suggesting an alteration in gamma frequency trace in the adapted condition, and such time-ranges are marked in Figure 2C and F by horizontal bars at the top.

### Jadi-Sejnowski (JS) model

Jadi and Sejnowski (Jadi and Sejnowski, 2014) used a simple rate model consisting of an excitatory and an inhibitory population with sigmoidal activation, operating as an ISN and constrained the input drives to the populations to reproduce the increase in power and decrease in peak frequency of gamma with increasing stimulus size as earlier studies have observed in V1 (Gieselmann and Thiele, 2008; Ray and Maunsell, 2011).

The model defines the population firing rates r_E_ and r_I_ of Excitatory and Inhibitory populations respectively as follows:

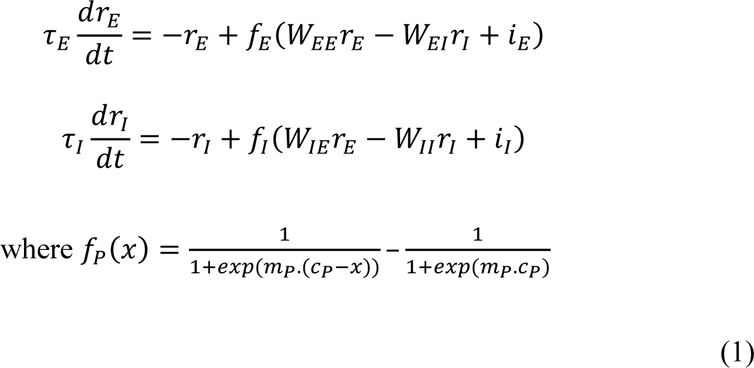

The JS model originally used constant input drives i_E_ and i_I_. In this model, larger stimuli cause an increased inhibitory drive (i_I_), owing to suppression from the surrounding populations. Since the model operated close to a supercritical Hopf bifurcation, the amplitude and frequency of oscillations in firing rates could be closely approximated by linearization of the model. The authors deduced and demonstrated that the model gave rise to the observed trends in the gamma when the inputs were such that the inhibitory population was strongly ‘superlinear’. This means that the summed inputs to the inhibitory population (from recurrent and external sources; argument of f_I_ in equation (1)) must lie in a certain range of values where the activation function σ_I_ curves upwards (increasing in slope with increasing summed input) (Jadi and Sejnowski, 2014). For the sigmoidal activation function used in equation (1), the summed inputs must operate in the lower half of the sigmoid. Superlinear activation of the excitatory population, on the other hand, was antagonistic (not strongly superlinear). The set of such inputs constitute the operating regime of the model, which we refer to as the ‘superlinear’ regime. The parameter combination used in this study is given in Table 1. The connectivity weights were retained as that of the original model by Jadi and Sejnowski (Jadi and Sejnowski, 2014) but c_E_ and c_I_ (θ_E_ and θ_I_ in Jadi and Sejnowski (2014)) were changed to ensure that our baseline period input combination (i_E_,i_I_) = (0,0) does not have multiple fixed points but yields steady-state value of firing rates close to 0. Further, the time-constants τ_E_ and τ_I_ were scaled by a factor of 0.64 from the original parameters to ensure that the model generates a wide range of limit cycles frequencies spanning the gamma range.

**Table 1:** Parameter values used in JS model with noisy inputs.

| $W_{EE}$ | $W_{EI}$ | $W_{IE}$ | $W_{II}$ | $\tau_E, \tau_I$<br>(ms) |
| --- | --- | --- | --- | --- |
| 16 | 26 | 20 | 1 | 12.8, 6.4 |
| $m_E, m_I$ | $c_E, c_I$ | $\theta_E, \theta_I$<br>(Hz) | $I_{E, min}, I_{I, min}$ | $I_{E, max}, I_{I, max}$ |
| 1, 1 | 9.5, 17 | 16, 1 | 2, 2 | 10.5, 7.5 |

In our study, we subject the model to noisy input drives i_E_ and i_I_, which were both Ornstein-Uhlenbeck (OU) noise (as in our previous study in Krishnakumaran and Ray 2023). The OU noise formulation of i_E_ and i_I_ was a first-order low pass filtered white Gaussian noise:

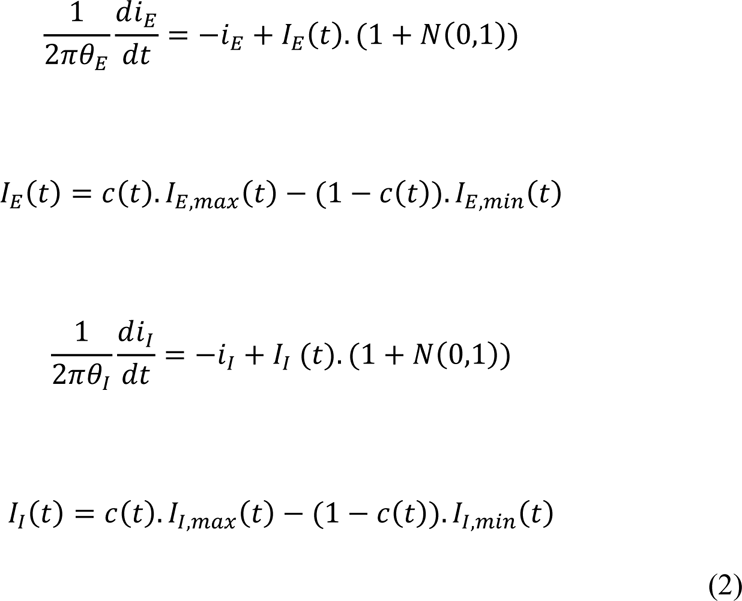

where I_E_ and I_I_ denote the desired steady-state mean values of i_E_ and i_I_ respectively, and *N*(0,1) denotes that at each timepoint the value was sampled from a Normal distribution. Values of I_E_ and I_I_ were determined by the contrast of the grating presented at time *t*, denoted as *c*(*t*). The contrast variable *c*(*t*) was varied between 0 and 1, indicating 0% and 100% contrasts respectively. For the contrast adaptation simulation, the lower contrast was emulated by setting *c*(*t*) as 0.66, while c(*t*)=1 was used as the adapting contrast. The experiment was simulated by presenting 3 contrasts values in sequence, namely c(*t*)=0.66, 1, 0.66, during intervals 0-1 s, 1-2 s and 2-3 s respectively. The first and second presentations of c(*t*)=0.66 were considered as the unadapted and the adapted conditions respectively.

The above formulation could be considered as an Additive White Gaussian (AWG) noise passed through a first-order low pass filter with a cutoff frequency at θ_P_ Hz for a population P. Further, at steady state, the variance would be π*I*_*P*_^2^θ_*P*_. Hence, θ_P_ could be assigned the unit Hz, whose lower value corresponds to the smoothly varying input drive to the population P with slow rise and fall, with less fluctuations and hence, less variance.

In the final simulation experiment, contrast of input was varied as a rectified sine profile as in Fig. 3 of Ray and Maunsell (2010). The contrast variation was simulated as:

**Figure 3:**
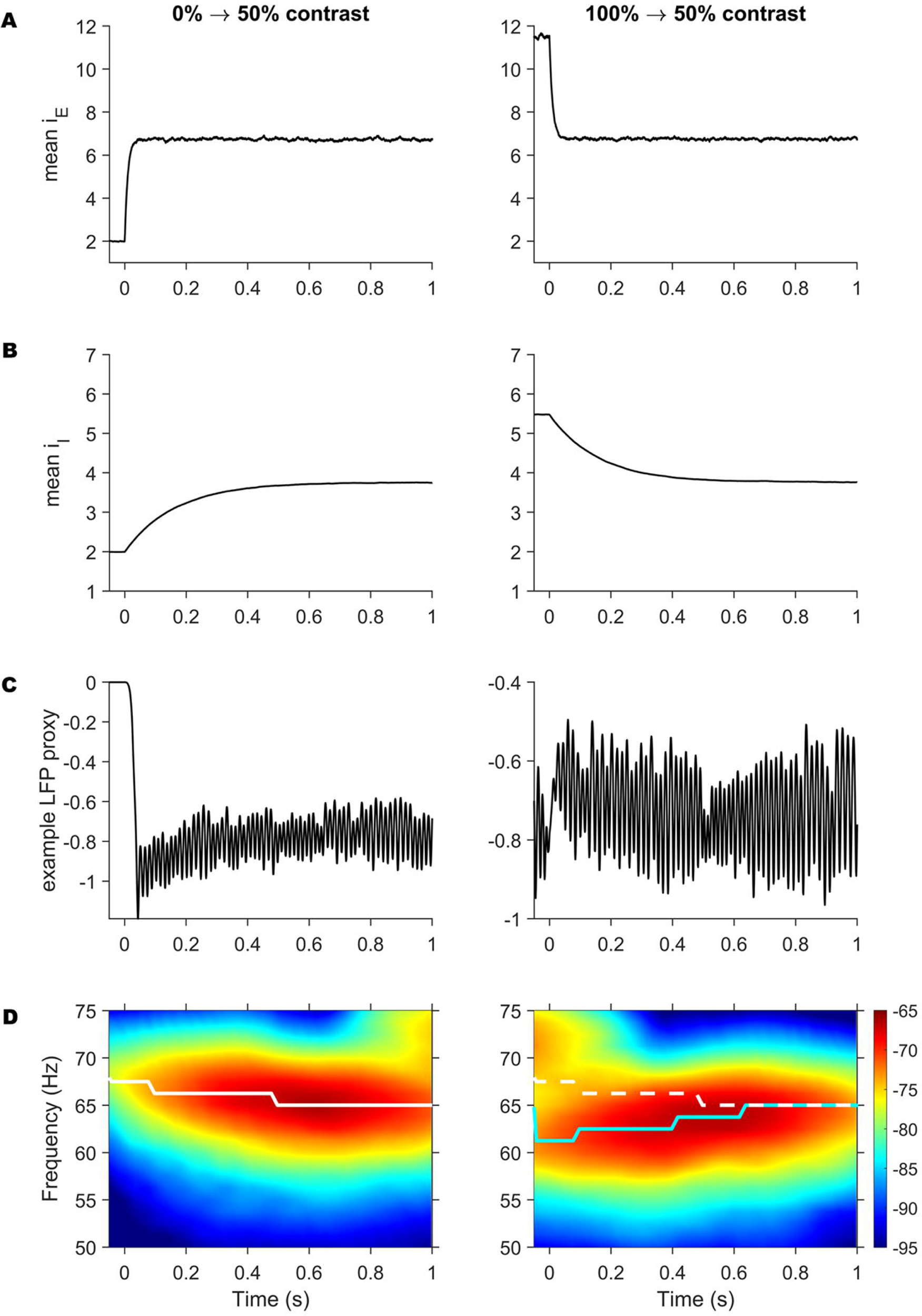
Gamma response in Noisy JS model with delayed inhibition to contrast adaptation. (A) Mean input drive to excitatory population (IE) estimated across 50 iterations for unadapted (left) and higher-contrast adapted (right) conditions. (B) Mean input drive inhibitory population (II) for the corresponding stimulus conditions. (C) LFP proxy output from an example iteration from the model in each condition. (D) Mean LFP time-frequency spectrum across iterations for each condition. White trace in both plots corresponds to the peak-frequency of the unadapted conditions. Cyan trace in right plot corresponds to the peak-frequency of the adapted condition.

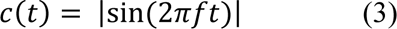

where the sinusoidal frequency f=0.625, 1.25 or 2.5 Hz. These stimuli were presented for 0.4 s duration while c(*t*)=0 for the rest of the simulation time.

### Spectral analysis of model output

Since the model was used with stochastic inputs, each input combination was simulated for 50 repeats. Each repeat spanned from -0.547 to 3.548 s and was simulated by the forward Euler method with a step size of 2e-5 s. An LFP proxy was computed from r_E_ and r_I_ as -r_E_-r_I_ to emulate the recording depth expected to be closer to apical dendrite distribution of L23 (as in our previous studies, (Krishnakumaran et al., 2022; Krishnakumaran and Ray, 2023); see Krishnakumaran et al. (2022) for more discussion, and Mazzoni et al. (2015) for a computational study of LFP proxies in a morphological network model).

The TF spectrum of the LFP proxy for the contrast-adaptation experiment was computed using MT in the same way as for the data, treating all 50 iterations of each stimulus condition as analogous to different trials from a single electrode.

However, for the rectified-sinusoidal contrast, the TF spectra were computed using Matching Pursuit (MP) (Mallat and Zhang, 1993). MP was used so that our simulated TF spectra can be compared against the MP TF spectra of LFP data in Fig. 3 of Ray and Maunsell (2010). To reduce computational load during MP, the LFP proxy signal recorded was low-pass filtered at a cutoff of 200 Hz using a 4^th^ order Butterworth filter and downsampled by 50, yielding 4096 datapoints. The resultant signal was subject to MP analysis with a dyadic dictionary, which iteratively fits gabors to the LFP traces and computes the TF spectrum of the traces by adding together the Wigner-Ville distribution of all these gabors (see Ray et al. (2008) and Chandran et al. (2016) for details of the algorithm and the merits of MP analysis over other methods).

## Results

LFP data was recorded from V1 in two awake macaques M1 and M2 in response to an fullscreen achromatic grating presented in different contrasts consecutively without interstimulus interval (as shown in Figure 1) from 77 and 32 electrodes, respectively. Figure 2 compares the change in TF spectra of LFP corresponding to a 50% contrast when preceded by a blank grey screen (unadapted) versus by a 100% contrast (adapted). The results of adaptation of 25% contrast were similar and are presented in Supplementary Figures S1 and S2 along with other contrast sequences.

### Gamma frequency transient vanishes after contrast adaptation

Figure 2A and D show the change in TF spectral power from baseline corresponding to the unadapted condition from M1 and M2 respectively. Shortly after onset of 50% contrast grating at 0 s, there was an increase in power at all frequencies corresponding to the onset response, followed by gamma rhythm starting at a high frequency and gradually decreasing in frequency to a steady-state value over the initial 500 ms. This is a classic “frequency-falloff” effect observed in many studies and modelled using slowly arriving inhibitory inputs in our previous study (Krishnakumaran and Ray 2023). Figure 2B and E show the change in TF spectra for the adapted condition with 100% contrast presented for 1 s before the 50% contrast. We hypothesized that in this case, both excitation and inhibition would be higher before the onset of the 50% contrast, and since the inhibition would take longer to dissipate, the frequency-falloff should be abolished or reversed. Consistent with this hypothesis, we indeed observed that the gamma band for the adapted condition did not exhibit a frequency-falloff in M1 (Figure 2B), and was even mildly reversed in M2 (Figure 2E). The peak-frequency trace of gamma was computed individually for each selected electrode, and shown as median±SE across electrodes in Figure 2C and F. Using a paired sign-test to verify this suppression of gamma frequency transient at each timepoint, we found p<0.01 for the entire duration in M1 and an initial ∼500 ms after the onset response in M2 after the onset response as indicated by the shaded areas at the top.

Supplementary Figures S1 and S2 show the change in TF spectra for M1 and M2 respectively for each pair of consecutive contrasts. For both 25% (top row) and 50% (middle row), increasing the contrast of the preceeding (adaptor) stimulus caused a reduction and subsequent reversal of the frequency fall-off effect in both monkeys (columns from left to right).

### Noisy JS model with delayed surround replicates contrast adaptation transients

In Figure 3, we replicated the above experiment in the noisy JS model with 50 iterations. In Figure 3A-B, mean input drive to excitation i_E_ and inhibition i_I_ respectively were estimated using equation (2), for the unadapted (left) and adapted (right) conditions. Due to the difference in θ between excitatory and inhibition inputs, the excitatory drive i_E_ had almost instantaneous onset/offset response, following the contrast variations, whereas the inhibitory drive i_I_ was slower and smoother with delayed response to stimulus onset. In Figure 3C, we show an example LFP proxy obtained from the noisy JS in one iteration, for each condition. After the lower contrast stimulus onset (marked as 0 s), in the unadapted condition (left), gamma rhythms were generated in bursts throughout the contrast presentation. Additionally, we found an initial evoked response in the baseline of the LFP proxy itself. In the unadapted condition, the non-zero stimulus onset evoked a negative deflection in the LFP proxy which disappears over time to settle at a steady-state value (higher baseline). However, in the adapted condition (right), the initial baseline deflection with respect to the later steady-state was in the opposite direction to that of the unadapted condition. In Figure 3D, the TF spectra averaged across 50 iterations of each condition are shown along with the peak frequency traces. We found an initial decrease in gamma frequency in the unadapted condition. However, in the adapted condition, the gamma frequency during the initial transient period was lower than the unadapted case. These differences in transients after adaptation were due to the slow-accumulating inhibitory inputs i_I_ as adaptation with higher contrast gave rise to a higher initial drive to inhibitory populations after onset of the 50% contrast than in the unadapted case (Figure 3B). In the unadapted condition, the slow increase of inhibitory input drive gave rise to the frequency-falloff, whereas in the adapted condition, the slow decay of inhibitory drive produced the opposite trend.

### Gamma frequency profile in response to smoothly varying contrast

In Figure 4, we present the output of the noisy JS model for rectified-sinusoidally time-varying contrasts to compare against the macaque LFP data from Ray and Maunsell (2010). In their experiment (see Fig. 3 of Ray and Maunsell (2010)), an achromatic grating was presented in time-varying contrast profile, where the profile varied as a rectified sine function of time emulated in equation (3), with frequencies f=0.625, 1.25 and 2.5 Hz. The resultant TF spectra showed time-varying gamma frequency with asymmetric rise and fall trends at higher frequencies, with sudden rises and gradual falls in frequency around a contrast peak. Figure 4A-B shows the mean excitatory and inhibitory drives emulating this experiment over 50 iterations. The excitatory drive traced the contrast variation closely, whereas the inhibitory drive rose and fell slowly resulting in delayed onset at lower frequencies and an accumulation across successive contrast crests at 2.5 Hz. Figure 4C shows example LFP proxies for the corresponding cases. For 0.625 Hz, although i_E_ increased early, the LFP proxy did not show any activity until it reached a higher value. This was due to the fact that the JS model shows 0 activity outside its oscillatory regime across the Hopf bifurcation for lower i_E_ (see Fig. 6 in Krishnakumaran and Ray (2023)). In our experiment, we operated the model between non-oscillating and oscillating inputs for lower and higher contrasts, respectively. As the contrast effects in the model were simulated by varying I_E_ more rapidly than I_I_, the model produced 0 response until I_E_ reached a higher value, leading to a delayed onset of the rhythm. On the other hand, at the falling phase of the contrast profile, since the inhibition delayed slowly, the rhythm was maintained for a longer duration with progressively decreasing frequency before going off the oscillatory regime. Figure 4D shows the TF spectra averaged across 50 iterations of each case. Gamma band increased in frequency slowly after the onset response for 0.625 Hz, and showed asymmetric frequency rise and fall around the contrast peak time for higher frequencies for 1.25 Hz, as seen in Fig. 3 of Ray and Maunsell (2010). In 2.5 Hz case, the decrease in i_E_ rapidly moved the model out of oscillatory regime, resulting in the gamma band being broken, which could also be seen in Fig. 3B of Ray and Maunsell (2010).

**Figure 4:**
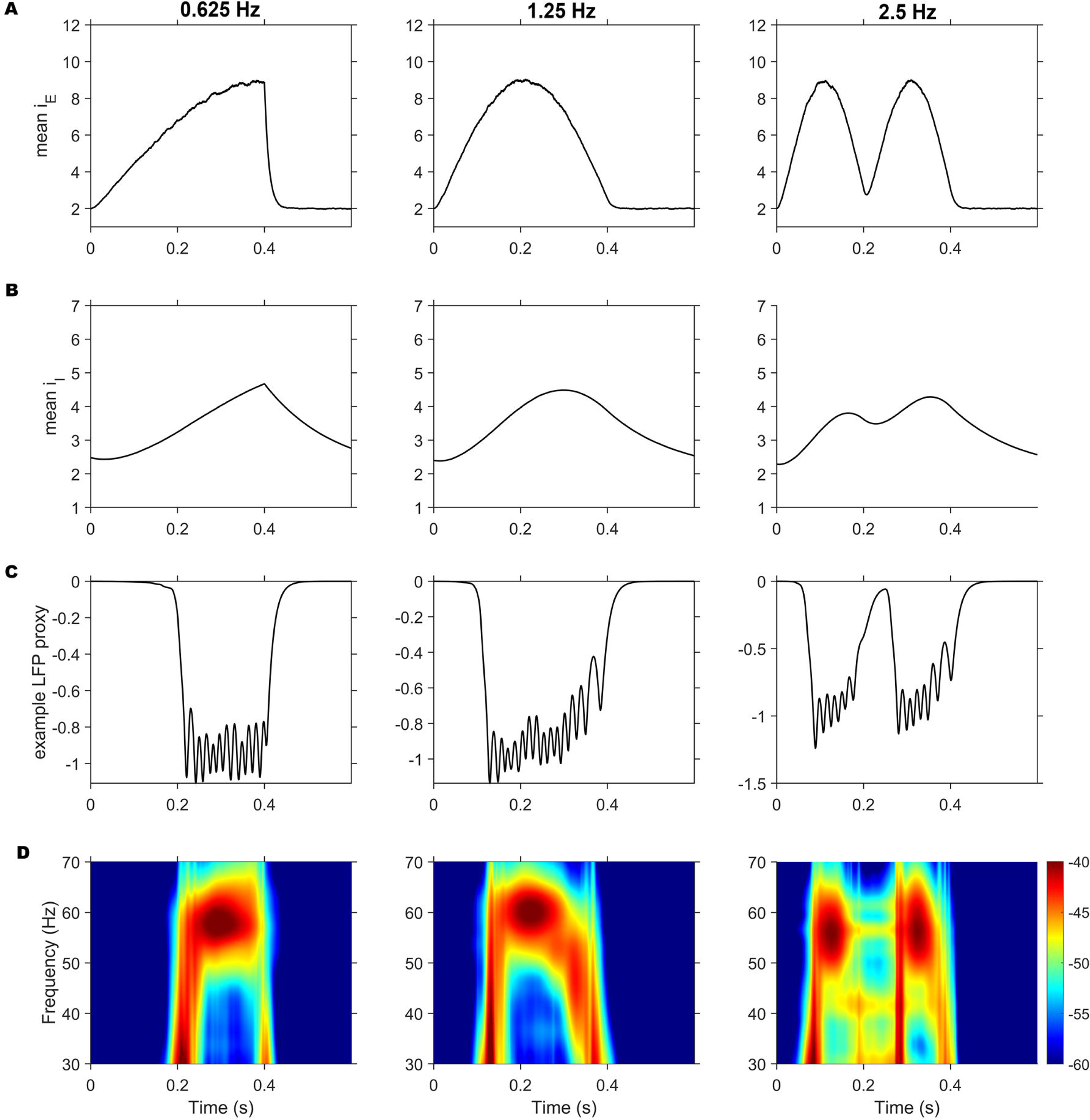
Gamma response in Noisy JS model with delayed inhibition to time-varying contrasts. (A) Mean input drive to excitatory population across 50 iterations for rectified sine contrast modulation of 0.625 Hz (left), 1.25 Hz (middle) and 2.5 Hz (right) conditions. (B) Mean input drive to inhibitory population for corresponding conditions. (C) LFP proxy output from an example iteration in each condition. (D) Mean LFP time-frequency spectrum across iterations in each condition.

## Discussion

We show that adaptation with a higher contrast grating abolishes or reverses the transient reduction in gamma frequency after onset of a grating. We show that this difference in gamma frequency transient after adaptation could be replicated in the noisy JS model by virtue of its delayed inhibitory input drive, which also replicates the variation in gamma onset and frequency modulation to stimuli with time-varying contrasts.

### Visual adaptation

Typical experiments for studying V1 circuitry involve sustained visual stimulation with constant, or ‘regularly’ varying stimuli, optimized for V1 receptive fields, whereas realistic stimulation and the corresponding responses are less parametrized but have been less tangible to inference so far. Stimulus-adaptation studies have shown that neuronal responses are dependent on the sequence of stimulus presentations, at every stage of the visual pathway from retinal ganglia to V1 (see Kohn (2007) for a review), and exhibit correlates to perceptual after-effects (Jin et al., 2005). Prior-dependent response to subsequent stimuli is ubiquitous in the sensory cortex, resulting in theories, such as predictive-coding or energy efficient-coding strategies (Webster, 2015; Weber et al., 2019), treating adaptation as a signature of the underlying perceptual-coding principle.

In V1, adaptation to prior exposure to achromatic stimuli (adaptors) reportedly result in temporary changes to orientation tuning for subsequent stimuli in the neurons recorded (Patterson et al., 2013; Ghodrati et al., 2019). The extent of these changes themselves vary in a graded fashion depending on the orientation of the adaptor-stimulus, suggesting a significant contribution from within V1 (Wissig and Kohn, 2012). Indeed, contributions to surround-suppression coming from lateral unmyelinated connections are orientation-selective (Angelucci et al., 2017), which is emulated in inhibitory drive dynamics in our model (see section *Delayed surround drive to inhibition* for elaboration). While the feedforward inputs to V1 from LGN are bound to exhibit temporary suppressive dynamics due to contrast adaptation themselves (Camp et al., 2009; Raghavan et al., 2023), in our model, we do not explicitly introduce these adaptation effects in the LGN-driven input drive I_E_ which may give rise to additional variations. Instead, we find that dynamics inherent to intra-cortical surround-suppression could describe the gamma frequency responses we observe in the LFP from V1.

### Delayed surround drive to inhibition

The noisy JS model replicates gamma frequency trends by setting θ_I_ << θ_E_ in the input model, as the lower value of θ_I_ delays the change in surround drive to inhibition. Delayed inhibition over millisecond timescales is observed in cortical activity and emerges intrinsically in PING models (see Krishnakumaran and Ray (2023), *Discussion* section for review). Typically, a local neuronal population as in the noisy JS model is thought to be strongly “local” in interconnectivity (within a cortical column), but they may send lateral inputs to similarly-tuned surround populations. Such lateral inputs are more convergent onto inhibitory neurons, mediating surround suppression (Shushruth et al., 2012). Further, these surround inputs arriving through unmyelinated axons, are expected to have a considerable latency depending on distance between source and target populations. Bair and colleagues (Bair et al., 2003) analyzed surround suppression in macaque V1, produced by far and near surround stimuli, and found two distinct suppression components with different timescales - a slower component thought to arise through lateral connections at latencies upto ∼20 ms and a fast component arriving from feedback from higher areas with much less delay (see Fig 7A in Bair et al. (2003); See Angelucci et al. (2017) for detailed review of different surround inputs). Therefore, while the incoming excitation to a particular area responds with lower latency to the feedforward projection from the immediately upstream area (LGN in case of V1 neurons), the inhibition which is activated by the local and widespread surround network with distance-dependent latencies (Girard et al., 2001), will be both delayed as well as smoothed out. Stimulus-selective surround suppression indeed exhibits delayed onset (∼10 ms after evoked response onset) in LFP recordings (Angelucci et al., 2017; Wang et al., 2020). Such slow dynamics of inhibition also yield transients in non-oscillatory activity. In recent studies (Zhou et al., 2019, 2023), a delayed normalization (DN) model has been fitted to data from intra-cranial EEG under different stimulus conditions. The DN model has a different structure than the noisy JS model, but also incorporates relative delays in excitation and surround-driven inhibition using different time-constants. In these studies, fitting data to DN model yielded a surround time-constant of over 2x the excitatory time-constant, arguing for delayed effect of surround suppression contributing to the transient trends in non-oscillatory activity.

### Inhibition network in V1

Interneurons in V1 are thought to be driven mostly by intra-cortical excitatory populations than by feedforward inputs to V1. This excitation-driven inhibition is evidenced by different surround-modulation and transient properties (Ozeki et al., 2009; Shushruth et al., 2012) typically observed in recordings from cats and macaques, and serves as the principle behind all variations of PING models (Buzsáki and Wang, 2012), and ISN models (Tsodyks et al., 1997; Jadi and Sejnowski, 2014) such as the JS model used in this study. Most network models simulated gamma rhythms generated by interactions between homogeneous populations of excitatory and inhibitory neurons (Chariker et al., 2018; Zachariou et al., 2021). However, recent studies in mice have shown distinct interactions of different interneuronal sub-types (Adesnik et al., 2012; Veit et al., 2017). Network models incorporating these details may help study their roles in cortical processing (Wagatsuma et al., 2023). Two major interneuronal networks involve Parvalbumin (PV)-expressing neurons and Somatostatin (SOM) and Vasoactive Intestinal peptide (VIP)-expressing neurons. The PV and SOM-VIP neuronal networks differ in spatial-connectivity ranges (Adesnik et al., 2012), with SOM neuron activities showing longer-range spatial integration than PV. Further, while PV neurons synapse onto a pyramidal cell near the soma, SOM neurons’ synapses are found at more distal regions. The synapse location difference has been demonstrated, by morphological modelling (Headley et al., 2024), to affect the nature and extent of the interneurons’ participation during rhythmic activity of different frequencies. Further the wider-ranged SOM network could be implicated in emergence of a second, slow-gamma of lower frequency range, with different orientation selectivity (see Fig. 1 in Murty et al. (2018)) and a wider spatial coherence profile than previously observed fast gamma, during larger-size stimulus presentations (Murty et al., 2018). While our current model does not generate slow-gamma, it has been modelled either using a second inhibitory population with different dynamics (Keeley et al., 2017) or simulating an altered effective connectivity caused by larger stimuli activating farther networks interacting with the local network by wider horizontal connections (Han et al., 2021).

### Notes on the JS model

JS model was chosen for its self-oscillating behavior across a supercritical Hopf bifurcation, exhibiting V1 gamma-like trends in power and frequency. The model represents a local population of strongly-interconnected excitatory and inhibitory neurons with similar stimulus selectivity. The strong-recurrent excitatory causes the model to operate in ISN regime (Tsodyks et al., 1997; Jadi and Sejnowski, 2014). It is expected that such a local population be uniformly-interconnected so that a mean-field approximation could be made under uniform stimulus conditions. Larger stimuli evoke activity in surrounding populations, which also drive the local population along with the LGN feedforward-input from the stimulus centre. This surround-input is expected to be stronger for the wide-connecting inhibitory neurons than the strongly locally-recurrent excitatory neurons. Hence, JS model emulates larger-size stimuli using higher values of input drive to inhibition I_I_ (Jadi and Sejnowski, 2014). The bifurcation property and its sensitivity to recurrent-connectivity parameters also allows it to emulate the attenuation of gamma power by different discontinuities in the fullscreen grating (Shirhatti et al., 2022). Shirhatti and colleagues modelled discontinuity effects as a change in recurrent-connectivity among the activated local population. This could result from either some neurons becoming inactive due to the discontinuity or from the segment of discontinuity recruiting a different set of neurons at different spatial locations which have less interconnectivity than the population activated by a uniform grating. The resultant decrease in recurrent-connectivity was shown to shrink the bifurcation boundary enclosing the oscillation-generating inputs. Hence a certain input would be out of the oscillatory-regime in the discontinuous condition. Further, the non-linearity of the model produces non-sinusoidal gamma waveforms. In Krishnakumaran et al. (2022), we found that the waveform shape found in LFP could be replicated by a subset of oscillatory inputs by the JS model.

However, as inputs in the original model were constant over time, the model would give rise to sustained gamma oscillations as opposed to the bursty-gamma observed in LFP. Furthermore, response of the model to stimulus change was rapid and lacked the transients observed in data. Gamma bursts and transient trends in frequency have been replicated by driving the model with OU-type noise in Krishnakumaran and Ray (2023) and in this study. While the core model and its interconnectivity has not been changed, the rich temporal properties of gamma rhythms emerge naturally when realistic input dynamics are incorporated. Still, the model’s specific non-linearity poses a hindrance to modelling realistic gradual-decrease in activity expected when contrast is slowly increased, as observed in Figure 4C for 0.625 Hz stimuli. Also, between non-oscillating and oscillatory values of I_E_, the model produces hysteresis over a set of I_E_ values with two stable fixed point, corresponding to different extreme steady-state population activities for each input, depending on prior activity. This, along with noisy inputs, results in deviating trajectories in some trials, contributing to the irregular low-power bursts of oscillation in the TF spectrum in Figure 4D for 2.5 Hz in the interval -0.2-0.3ms. Overcoming these shortcomings would require abandoning the existing, simple non-linearity of the model to accommodate realistic behavior outside the oscillation-generating input domain. However, such a data-driven alteration of the model’s non-linearity has not yet been attempted.

## Acknowledgements

We thank Surya S Prakash, Ankan Biswas and Divya Gulati for their assistance in data collection. We thank Prof. Adam Kohn for valuable discussions and insights.

## Supplementary Figures

**Figure S1:**
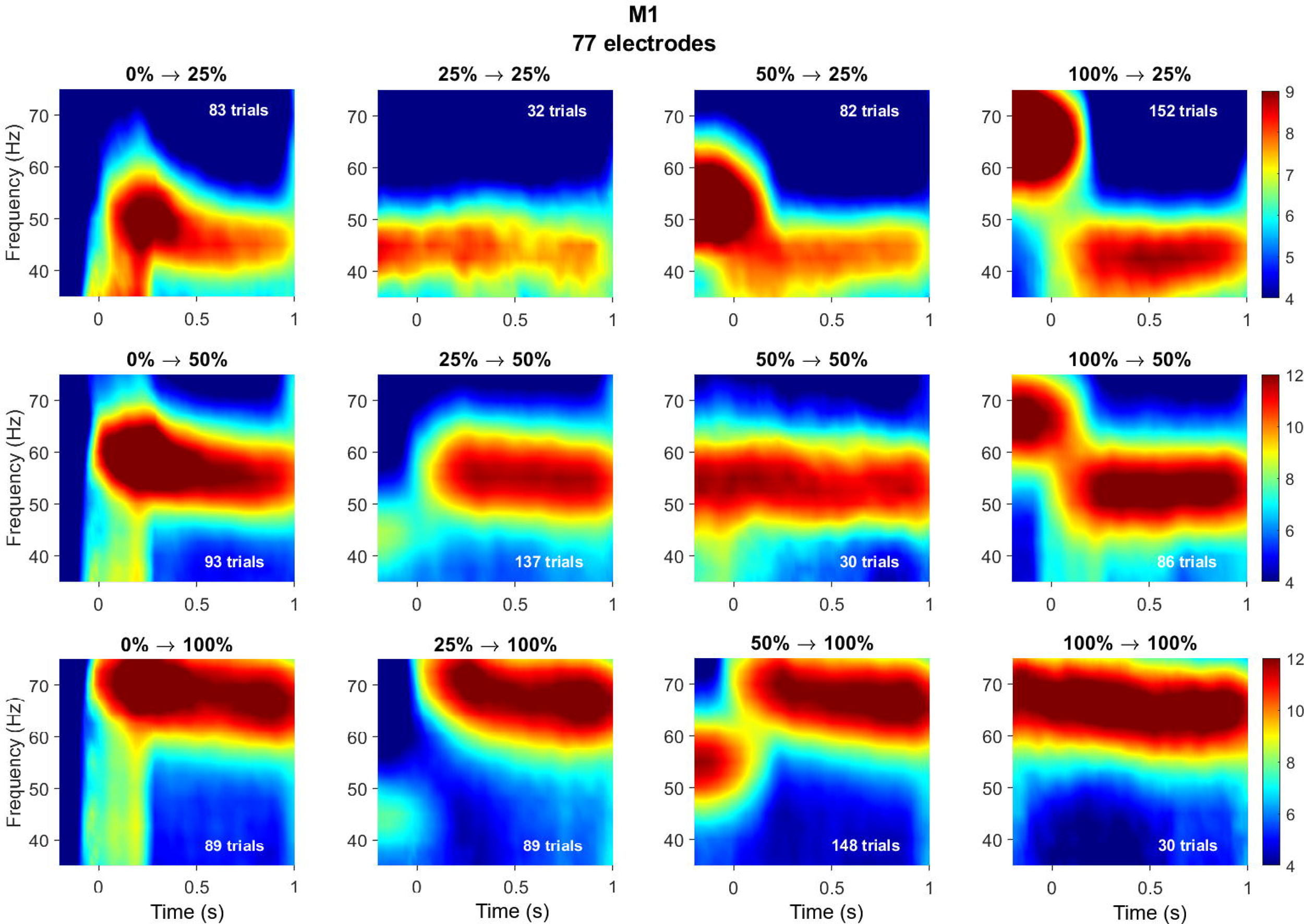
Change in time-frequency spectra for different consecutively presented contrast pairs in M1. Each column corresponds to a different ‘adaptor’ contrast. Each row corresponds to a different ‘adapted’ contrast, that follows the adaptor. The sequence of contrasts is indicated in text above the respective plots. The plots corresponding to trials where the same contrast was presented for >2 s continuously occupy a diagonal of the figure. Note that these same contrast trials are much lower compared to other conditions, due to which the TF spectra are very patchy. For each contrast pair, the number of good trials used for TF analysis are indicated in text within the respective plot. Colorbars for each row are given at the right of the respective row. Note that colorbars have different scales across rows for better visualization of gamma frequency variation.

**Figure S2:**
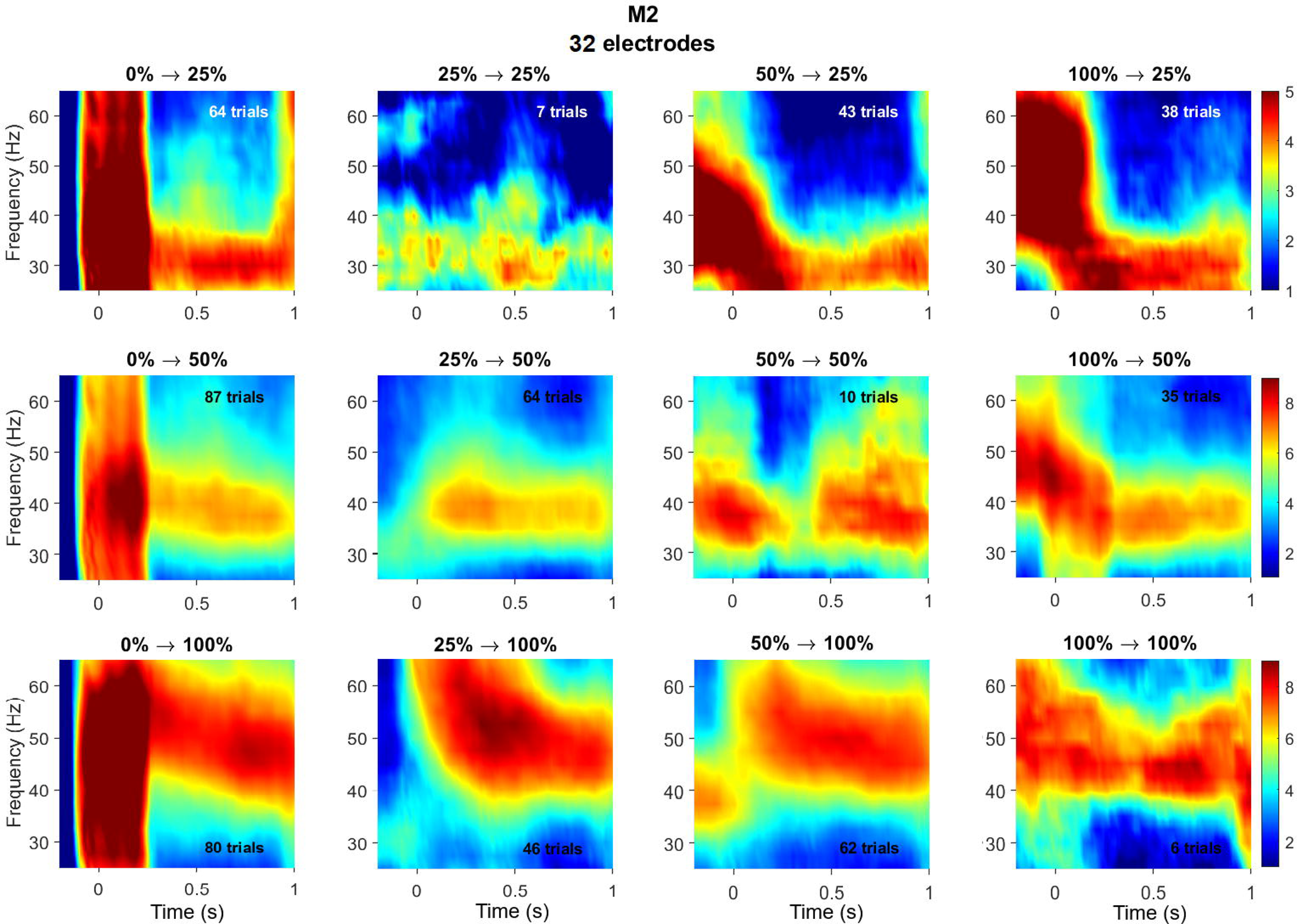
Change in time-frequency spectra for different consecutively presented contrast pairs in M2. Same as Figure S1, but for M2. Patchy TF spectra seen for prolonged presentation of the same contrast are due to less number of such trials.

## References

Adesnik H, Bruns W, Taniguchi H, Huang ZJ, Scanziani M. 2012. A neural circuit for spatial summation in visual cortex. Nature 490:226–231.

Angelucci A, Bijanzadeh M, Nurminen L, Federer F, Merlin S, Bressloff PC. 2017. Circuits and Mechanisms for Surround Modulation in Visual Cortex. Annual Review of Neuroscience 40:425–451.

Bair W, Cavanaugh JR, Movshon JA. 2003. Time Course and Time-Distance Relationships for Surround Suppression in Macaque V1 Neurons. J Neurosci 23:7690–7701.

Bokil H, Andrews P, Kulkarni JE, Mehta S, Mitra PP. 2010. Chronux: A platform for analyzing neural signals. Journal of Neuroscience Methods 192:146–151.

Buzsáki G, Wang X-J. 2012. Mechanisms of Gamma Oscillations. Annual Review of Neuroscience 35:203–225.

Camp AJ, Tailby C, Solomon SG. 2009. Adaptable Mechanisms That Regulate the Contrast Response of Neurons in the Primate Lateral Geniculate Nucleus. J Neurosci 29:5009– 5021.

Chariker L, Shapley R, Young L-S. 2018. Rhythm and Synchrony in a Cortical Network Model. J Neurosci 38:8621–8634.

Dubey A, Ray S. 2019. Cortical Electrocorticogram (ECoG) Is a Local Signal. J Neurosci 39:4299–4311.

Ghodrati M, Zavitz E, Rosa MGP, Price NSC. 2019. Contrast and luminance adaptation alter neuronal coding and perception of stimulus orientation. Nat Commun 10:941.

Gieselmann MA, Thiele A. 2008. Comparison of spatial integration and surround suppression characteristics in spiking activity and the local field potential in macaque V1. Eur J Neurosci 28:447–459.

Girard P, Hupé JM, Bullier J. 2001. Feedforward and Feedback Connections Between Areas V1 and V2 of the Monkey Have Similar Rapid Conduction Velocities. Journal of Neurophysiology 85:1328–1331.

Han C, Wang T, Wu Y, Li Y, Yang Y, Li L, Wang Y, Xing D. 2021. The Generation and Modulation of Distinct Gamma Oscillations with Local, Horizontal, and Feedback Connections in the Primary Visual Cortex: A Model Study on Large-Scale Networks. Neural Plasticity 2021:e8874516.

Headley DB, Latimer B, Aberbach A, Nair SS. 2024. Spatially targeted inhibitory rhythms differentially affect neuronal integration. eLife 13 Available at: https://elifesciences.org/reviewed-preprints/95562 [Accessed June 17, 2024].

Jadi MP, Sejnowski TJ. 2014. Regulating Cortical Oscillations in an Inhibition-Stabilized Network. Proc IEEE Inst Electr Electron Eng 102.

Jin DZ, Dragoi V, Sur M, Seung HS. 2005. Tilt Aftereffect and Adaptation-Induced Changes in Orientation Tuning in Visual Cortex. Journal of Neurophysiology 94:4038–4050.

Keeley S, Fenton AA, Rinzel J. 2017. Modeling fast and slow gamma oscillations with interneurons of different subtype. Journal of neurophysiology 117:950–965.

Kohn A. 2007. Visual Adaptation: Physiology, Mechanisms, and Functional Benefits. Journal of Neurophysiology 97:3155–3164.

Krishnakumaran R, Raees M, Ray S. 2022. Shape analysis of gamma rhythm supports a superlinear inhibitory regime in an inhibition-stabilized network. PLOS Computational Biology 18:e1009886.

Krishnakumaran R, Ray S. 2023. Temporal characteristics of gamma rhythm constrain properties of noise in an inhibition-stabilized network model. Cerebral Cortex 33:10108–10121.

Mallat SG, Zhang Z. 1993. Matching pursuits with time-frequency dictionaries. IEEE Transactions on Signal Processing 41:3397–3415.

Mazzoni A, Lindén H, Cuntz H, Lansner A, Panzeri S, Einevoll GT. 2015. Computing the Local Field Potential (LFP) from Integrate-and-Fire Network Models. PLOS Computational Biology 11:e1004584.

Murty DV, Manikandan K, Kumar WS, Ramesh RG, Purokayastha S, Nagendra B, ML A, Balakrishnan A, Javali M, Rao NP, Ray S. 2021. Stimulus-induced gamma rhythms are weaker in human elderly with mild cognitive impairment and Alzheimer’s disease Vinck M, Colgin LL, Bosman CA, eds. eLife 10:e61666.

Murty DVPS, Manikandan K, Kumar WS, Ramesh RG, Purokayastha S, Javali M, Rao NP, Ray S. 2020. Gamma oscillations weaken with age in healthy elderly in human EEG. NeuroImage 215:116826.

Murty DVPS, Shirhatti V, Ravishankar P, Ray S. 2018. Large Visual Stimuli Induce Two Distinct Gamma Oscillations in Primate Visual Cortex. J Neurosci 38:2730–2744.

Ozeki H, Finn IM, Schaffer ES, Miller KD, Ferster D. 2009. Inhibitory stabilization of the cortical network underlies visual surround suppression. Neuron 62:578–592.

Patterson CA, Wissig SC, Kohn A. 2013. Distinct Effects of Brief and Prolonged Adaptation on Orientation Tuning in Primary Visual Cortex. J Neurosci 33:532–543.

Perry G, Taylor NW, Bothwell PCH, Milbourn CC, Powell G, Singh KD. 2020. The gamma response to colour hue in humans: Evidence from MEG. PLOS ONE 15:e0243237.

Peter A, Uran C, Klon-Lipok J, Roese R, van Stijn S, Barnes W, Dowdall JR, Singer W, Fries P, Vinck M. 2019. Surface color and predictability determine contextual modulation of V1 firing and gamma oscillations Colgin L, ed. eLife 8:e42101.

Raghavan RT, Kelly JG, Hasse JM, Levy PG, Hawken MJ, Movshon JA. 2023. Contrast and Luminance Gain Control in the Macaque’s Lateral Geniculate Nucleus. eNeuro 10 Available at: https://www.eneuro.org/content/10/3/ENEURO.0515-22.2023 [Accessed June 20, 2024].

Ray S, Hsiao SS, Crone NE, Franaszczuk PJ, Niebur E. 2008. Effect of Stimulus Intensity on the Spike–Local Field Potential Relationship in the Secondary Somatosensory Cortex. J Neurosci 28:7334–7343.

Ray S, Maunsell JH. 2011. Different origins of gamma rhythm and high-gamma activity in macaque visual cortex. PLoS biology 9.

Ray S, Maunsell JHR. 2010. Differences in Gamma Frequencies across Visual Cortex Restrict Their Possible Use in Computation. Neuron 67:885–896.

Shirhatti V, Ravishankar P, Ray S. 2022. Gamma oscillations in primate primary visual cortex are severely attenuated by small stimulus discontinuities. PLOS Biology 20:e3001666.

Shushruth S, Mangapathy P, Ichida JM, Bressloff PC, Schwabe L, Angelucci A. 2012. Strong Recurrent Networks Compute the Orientation Tuning of Surround Modulation in the Primate Primary Visual Cortex. J Neurosci 32:308–321.

Tsodyks MV, Skaggs WE, Sejnowski TJ, McNaughton BL. 1997. Paradoxical effects of external modulation of inhibitory interneurons. J Neurosci 17:4382–4388.

Veit J, Hakim R, Jadi MP, Sejnowski TJ, Adesnik H. 2017. Cortical gamma band synchronization through somatostatin interneurons. Nat Neurosci 20:951–959.

Wagatsuma N, Nobukawa S, Fukai T. 2023. A microcircuit model involving parvalbumin, somatostatin, and vasoactive intestinal polypeptide inhibitory interneurons for the modulation of neuronal oscillation during visual processing. Cerebral Cortex 33:4459–4477.

Wang T, Li Y, Yang G, Dai W, Yang Y, Han C, Wang X, Zhang Y, Xing D. 2020. Laminar Subnetworks of Response Suppression in Macaque Primary Visual Cortex. J Neurosci 40:7436–7450.

Weber AI, Krishnamurthy K, Fairhall AL. 2019. Coding Principles in Adaptation. Annual Review of Vision Science 5:427–449.

Webster MA. 2015. Visual Adaptation. Annual Review of Vision Science 1:547–567.

Wissig SC, Kohn A. 2012. The influence of surround suppression on adaptation effects in primary visual cortex. J Neurophysiol 107:3370–3384.

Xing D, Shen Y, Burns S, Yeh C-I, Shapley R, Li W. 2012. Stochastic Generation of Gamma-Band Activity in Primary Visual Cortex of Awake and Anesthetized Monkeys. J Neurosci 32:13873–13880a.

Zachariou M, Roberts MJ, Lowet E, De Weerd P, Hadjipapas A. 2021. Empirically constrained network models for contrast-dependent modulation of gamma rhythm in V1. NeuroImage 229:117748.

Zhou J, Benson NC, Kay K, Winawer J. 2019. Predicting neuronal dynamics with a delayed gain control model. PLOS Computational Biology 15:e1007484.

Zhou J, Whitmire M, Chen Y, Seidemann E. 2023. Delayed normalization model captures disparate nonlinear neural dynamics measured with different techniques in macaque and human V1. :2023.01.30.525700 Available at: https://www.biorxiv.org/content/10.1101/2023.01.30.525700v1 [Accessed June 10, 2024].

